# Gamma oscillations weaken with age in healthy elderly in human EEG

**DOI:** 10.1101/696781

**Authors:** Dinavahi V. P. S. Murty, Keerthana Manikandan, Wupadrasta Santosh Kumar, Ranjini Garani Ramesh, Simran Purokayastha, Mahendra Javali, Naren Prahalada Rao, Supratim Ray

## Abstract

Gamma rhythms (∼20-70 Hz) have been reported to be abnormal in mental disorders such as autism and schizophrenia in humans, and Alzheimer’s disease (AD) models in rodents. However, the effect of normal aging on these oscillations is not known, especially for elderly subjects (>49 years) for which AD is most prevalent. In a first large-scale (236 subjects; 104 females) electroencephalogram (EEG) study on gamma oscillations on elderly subjects (aged 50-88 years), we presented full-screen Cartesian gratings that induced two distinct gamma oscillations (slow: 20-34 Hz and fast: 36-66 Hz). Power and centre frequency significantly decreased with age for both slow and fast gamma, but not alpha (8-12 Hz). Reduction was more salient for fast gamma than slow. These results were independent of microsaccades and pupillary reactivity to stimulus, as well as variations in power spectral density with age. Steady-state visual evoked potentials (SSVEPs) also reduced with age. These results are crucial first steps towards using gamma/SSVEPs as biomarkers of cognitive decline in elderly.

**Significance statement:** No study in humans has examined visual narrow-band gamma oscillations with healthy aging in elderly subjects. In a first large-scale (236 subjects) EEG study on stimulus-induced narrow band gamma in cognitively normal elderly (>49 years) humans, we show that both power and centre frequency of slow and fast gamma (20-34 Hz and 36-66 Hz respectively) decrease with age, but alpha power does not. Steady-state visual evoked potential (SSVEPs) in gamma range also decrease with age. Any EEG-based biomarker is accessible and affordable to patients of a wide socio-economic spectrum. Our results are important steps for developing such screening/diagnostic tests for aging-related diseases, like Alzheimer’s disease.

## Introduction

Gamma rhythms are narrow-band oscillations often observed in the electrical activity of the brain, with centre frequency occupying ∼20-70 Hz frequency range. Previous studies have proposed involvement of these rhythms in certain higher cognitive functions like feature binding (Gray et al., 1989), attention (Gregoriou et al., 2009; Chalk et al., 2010) and working memory (Pesaran et al., 2002). Further, some studies have shown that these rhythms may be abnormal in neuropsychiatric disorders such as schizophrenia (Tada et al., 2014; Hirano et al., 2015), autism (Uhlhaas and Singer, 2007; Wilson et al., 2007; An et al., 2018) and Alzheimer’s disease (AD; Palop and Mucke, 2016; Mably and Colgin, 2018).

Gamma rhythms can be induced in the occipital areas by presenting appropriate visual stimuli such as bars and gratings, and their magnitude and centre frequency critically depend on the properties of the stimulus such as contrast, size, orientation, spatial frequency and drift rate (Jia et al., 2013; Ray and Maunsell, 2015; Murty et al., 2018). Recently, we showed that large (full-screen) gratings induce two distinct narrow-band gamma oscillations in local field potentials (LFP) in macaque V1 and posterior electrodes in human EEG, which we termed slow (∼20-40 Hz) and fast (∼40-70 Hz) gamma (Murty et al., 2018). Fast gamma was not a harmonic of slow, but instead these rhythms were differently tuned to stimulus properties. Importantly, slow gamma was observed only when the grating size was sufficiently large (diameter >8° of visual angle for humans). Two distinct gamma rhythms have also been recently reported in human MEG (Pantazis et al., 2018) and in visual cortex (Veit et al., 2017) and hippocampus (Colgin et al., 2009) in rodents. These rhythms have been suggested to be generated from excitatory-inhibitory interactions of pyramidal cell and interneuron networks (Buzsáki and Wang, 2012), specifically involving parvalbumin (PV) and somatostatin (SOM) interneurons (Cardin et al., 2009; Sohal et al., 2009; Veit et al., 2017).

A recent study has reported PV interneuron dysfunction in parietal cortex of AD patients and transgenic models of mice (Verret et al., 2012); and aberrant gamma activity in parietal cortex in such mice. However, our knowledge about these rhythms in healthy aging in humans is limited. Studies in human MEG have observed that the centre frequency of gamma oscillations is negatively correlated with age of healthy subjects in the range of 8-45 years (Muthukumaraswamy et al., 2010; Gaetz et al., 2012), but our understanding of these rhythms in elderly humans (>49 years), which is clinically more relevant for studying diseases of abnormal aging like AD, is lacking.

Further, visual stimulation of wild type and AD models of mice using light flickering at 40 Hz rescued AD pathology in visual cortex (Iaccarino et al., 2016). Such stimulation is known to entrain brain rhythms and generate steady-state visual evoked potentials (SSVEPs) at 40 Hz. However, to our knowledge, no previous study has examined SSVEPs in gamma band in healthy elderly. Furthermore, a recent study has shown flattening of power spectral density (PSD) in 2-24 Hz range in elderly subjects compared to younger subjects (Voytek et. al., 2015). However, how this flattening affects gamma rhythms in elderly has not been examined.

In this study, we described the variation of the two gamma rhythms in healthy elderly subjects. We first used a battery of cognitive tests to identify a large cohort (236 subjects; 104 females) of cognitively healthy elderly subjects aged between 50-88 years. For comparison, we also included 47 younger subjects (aged 20-48 years, 16 females). We induced gamma oscillations using full-screen static Cartesian gratings while monitoring their eye movements and recording EEG, and studied how slow and fast gamma and alpha oscillations, as well as slope of the PSD, varied with age in elderly subjects. In a subset of subjects, we examined SSVEPs in gamma frequency range (32 Hz) as well.

## Materials and Methods

### Human subjects

We recruited 236 elderly subjects (104 females) aged 50-88 years from the Tata Longitudinal Study of Aging cohort from urban communities in Bangalore through awareness talks on healthy aging and dementia. Recruitment was done by trained psychologists, who also collected their demographic details. Psychiatrists and neurologists at National Institute of Mental Health and Neurosciences (NIMHANS), Bangalore and M S Ramaiah Hospital, Bangalore assessed the cognitive function of these subjects using a combination of Clinical Dementia Rating scale (CDR), Addenbrook’s Cognitive Examination-III (ACE-III), Hindi Mental State Examination (HMSE), and other structured and semi-structured interviews. We considered only those subjects who were labelled as cognitively healthy for this study. Out of 236 cognitively healthy subjects thus recruited, we discarded data of 9 subjects (3 females) due to noise (see Artifact Rejection section below for details). We were thus left with 227 subjects (101 females) aged 50-88 years (mean±SD: 66.8±8.2 years) for analysis.

Further, we also recruited 47 younger subjects (16 females) aged 20-48 years (mean±SD: 30.4±7.1 years) from the student and staff community of Indian Institute of Science. We screened them orally for any past history of neurological/psychiatric illness. We had presented data from 10 of these younger subjects in an earlier study (Murty et al., 2018).

All subjects participated in the study voluntarily and against monetary compensation. We obtained informed consent from all subjects for performing the experiment. The Institute Human Ethics Committees of Indian Institute of Science, NIMHANS, Bangalore and M S Ramaiah Hospital, Bangalore approved all procedures.

### EEG recordings

Experimental setup, EEG recordings and analysis were similar to what we had described in our previous study (Murty et al., 2018). We recorded raw EEG signals from 64 active electrodes (actiCAP) using BrainAmp DC EEG acquisition system (Brain Products GmbH). We placed the electrodes according to the international 10-10 system. We filtered raw signals online between 0.016 Hz (first-order filter) and 1000 Hz (fifth-order Butterworth filter), sampled at 2500 Hz and digitized at 16-bit resolution (0.1 μV/bit). We rejected electrodes whose impedance was more than 25 KΩ. This led to a rejection of 3.9% of electrodes in elderly age-group (1.1% in younger subjects). However, most of these electrodes were frontal/central, and specifically, none were the ten parieto-occipital/occipital electrodes used for analyses (see Data Analysis subsection). Impedance of the final set of electrodes was 5.5±4.2 KΩ (mean±SD) for elderly subjects and 3.7±3.4 KΩ for younger subjects. We referenced EEG signals to FCz during acquisition (unipolar reference scheme).

### Experimental setting and behavioral task

All subjects sat in a dark room in front of an LCD screen with their head supported by a chin rest. The screen (BenQ XL2411) had a resolution of 1280 x 720 pixels and a refresh rate of 100 Hz. It was gamma-corrected and was placed at a mean±SD distance of 58.1±0.9 cm from the subjects (53.9-63.0 cm for all 274 subjects, 54.9-61.0 cm for the 227 elderly subjects) according to their convenience (thus subtending a width of at least 52° and height of at least 30° of visual field for full-screen gratings). We calibrated the stimuli to the viewing distance in all cases.

Subjects performed a visual fixation task. Stimulus presentation was done by a custom software running on MAC OS that also controlled the task flow. Every trial started with the onset of a fixation spot (0.1°) shown at the center of the screen, on which the subjects were instructed to hold and maintain fixation. After an initial blank period of 1000 ms, a series of stimuli (2 to 3) were randomly shown for 800 ms each with an inter-stimulus interval of 700 ms. Stimuli were sinusoidal luminance gratings presented full screen at full contrast. For the main “Gamma” experiment, these were presented at one of three spatial frequencies (SFs): 1, 2, and 4 cycles per degree (cpd) and one of four orientations: 0°, 45°, 90° and 135°. Subjects performed this task in 2-3 blocks (total 597 blocks across 283 subjects) during a single session according to their comfort. For an initial subset of subjects, stimuli with SF of 0.5 and/or 8 were also presented, but we discarded these SFs from further analysis to maintain uniformity. We also tested 32-Hz SSVEPs on a subset of the subjects who had analyzable data for the gamma experiment (221/227 elderly and 46/47 younger subjects) according to their willingness for the SSVEP experiment. Gratings (with SF and orientation that showed high change in slow and fast gamma power for each subject) counter-phased at 16 cycles per second (cps) in a similar stimulus presentation paradigm as described above, randomly interleaved with static gratings of the same SF and orientation.

### Eye position analysis

We recorded eye signals (pupil position and diameter data) using EyeLink 1000 (SR Research Ltd., sampled at 500 Hz) during the entire trial for all but one subject. We calibrated the eye-tracker for pupil position and monitor distance for each subject before the start of the session. All the subjects were able to maintain fixation with a standard deviation of less than 0.6° (elderly, eye-data for Gamma experiment shown in Figure 7a) and 0.2° (young, data not shown). We defined fixation breaks as eye-blinks or shifts in eye-position outside a square window of width 5° centered on the fixation spot. We rejected stimulus repeats with fixation breaks during −0.5s to 0.75s of stimulus onset, either online (and repeated the stimulus thus discarded), or offline (we took a few additional trials to compensate for possible offline rejection), according to the subjects’ comfort. This led to rejection of 16.7±14.2% (mean±SD) and 16.7±15.1% stimulus repeats for elderly subjects (for Gamma and SSVEP experiments respectively), most of who preferred offline rejection. For younger subjects, for many of whom we used online eye-monitoring, the rate of rejection due to fixation breaks was low (4.9±5.7% and 4.2±7.0%).

### Artifact rejection

We first estimated bad stimulus repeats for each unipolar electrode separately. We applied a trial-wise thresholding process on both raw waveforms (high-pass filtered at 1.6 Hz to eliminate slow trends if any) and multi-tapered PSD (computed between −500 ms to 750 ms of stimulus onset using the Chronux toolbox (Mitra and Bokil, 2008, http://chronux.org/, RRID:SCR_005547)). Any stimulus repeat for which either the waveform or the PSD deviated by 6 times the standard deviation from the mean at any time bin (between −500 ms to 750 ms) or frequency point (between 0-200 Hz) was considered a bad repeat for that electrode. We then created a common set of bad repeats across all 64 unipolar electrodes by first discarding those electrodes that had more than 30% of all repeats marked as bad, and subsequently assigning any repeat as bad if it occurred in more than 10% of total number of remaining electrodes. Finally, any repeat that was marked bad in any of the ten unipolar electrodes used for analysis (P3, P1, P2, P4, PO3, POz, PO4, O1, Oz, and O2; see Data Analysis subsection) was unconditionally included in the common bad repeats list, providing a final list of common bad repeats for each block for each subject. In spite of these stringent conditions, these led to a rejection of less than 20% of data (18.4±6.4% and 17.0±5.1% for elderly and younger subjects).

In addition, we calculated slopes (see Data Analysis subsection) of PSD (calculated with 1 taper and averaged across repeats, after removal of bad repeats) for each block in 56 Hz to 84 Hz range (to include the fast gamma range) for each unipolar electrode. Previous studies have shown that in clean electrophysiological data, PSD slopes are typically between 0.5 to 4.5 (Podvalny et al., 2015; Shirhatti et al., 2016; Muthukumaraswamy and Liley, 2018; Sheehan et al., 2018). We therefore discarded those electrodes (5.0±5.9% for elderly and 5.2±7.7% for younger subjects) that had PSD slopes less than 0. We further discarded any block (53/497 and 5/100 for elderly and younger subjects) that did not have at least a single clean bipolar electrode pair in any of the three groups of bipolar electrodes used for analysis (depicted in Figure 3d, see Data Analysis subsection for details): PO3-P1, PO3-P3, POz-PO3 (left anterolateral group); PO4-P2, PO4-P4, POz-PO4 (right anterolateral group) and Oz-POz, Oz-O1, Oz-O2 (posteromedial group). We then pooled data across all good blocks for every subject separately for final analysis. Those subjects who did not have any analyzable blocks (9/236 and 0/47 for elderly and younger subjects respectively) were discarded from further analysis, leaving 227 elderly (aged 50-88 years, mean±SD: 66.8±8.2 years, females: 101) and 47 young subjects (aged 20-48 years, mean±SD: 30.4±7.1 years, females: 16) for analysis. The total number of repeats per electrode that were finally analyzed were 276.2±87.2 for elderly subjects and 270.4±67.4 for younger subjects.

We applied a similar artifact rejection procedure for SSVEP experiment. Out of subjects with analyzable blocks for the Gamma experiment, 197 elderly (mean±SD: 66.8±7.8 years, females: 93) and 43 young (mean±SD: 30.4±7.3 years, females: 15) subjects had analyzable blocks (242/270) for SSVEP experiment. Using similar selection criteria as before, we rejected 7.7±5.2% of repeats for elderly subjects and 6.6±4.1% for younger subjects. The total number of analyzed repeats per electrode for counter-phasing condition were 30.2±6.9 and 29.7±6.6 for elderly and younger subjects respectively.

### EEG data analysis

For all analyses (unless otherwise mentioned), we used bipolar reference scheme. We re-referenced data at each electrode offline to its neighboring electrodes. We thus obtained 112 bipolar pairs out of 64 unipolar electrodes (Murty et al., 2018, depicted in Figure 3e). We considered the following bipolar combinations for analysis, except for scalp maps: PO3-P1, PO3-P3, POz-PO3 (left anterolateral group); PO4-P2, PO4-P4, POz-PO4 (right anterolateral group) and Oz-POz, Oz-O1, Oz-O2 (posteromedial group), depicted in Figure 3d. We discarded a bipolar electrode if either of its constituting unipolar electrodes was marked bad as described in the previous section. Data was pooled for the rest of the bipolar combinations in each of the electrode groups for further analysis.

We analyzed all data using custom codes written in MATLAB (The MathWorks, Inc, RRID:SCR_001622). We computed PSD and the time-frequency power spectrograms using multi-taper method with a single taper using Chronux toolbox. We chose baseline period between −500 ms to 0 ms of stimulus onset, while stimulus period between 250 ms to 750 ms to avoid stimulus-onset related transients, yielding a frequency resolution of 2 Hz for the PSDs. We calculated time frequency power spectra using a moving window of size 250 ms and step size of 25 ms, giving a frequency resolution of 4 Hz.

We calculated change in power in alpha rhythm and the two gamma rhythms as follows:

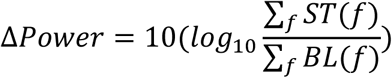

Where ST and BL are stimulus and baseline power spectra (across frequency *f*) averaged across repeats for all stimulus conditions and analyzable bipolar electrodes. For slow gamma, *f* ∈ [20 34] Hz and for fast gamma, *f* ∈ [36 66] Hz. Note that due to this averaging, on average more than 250 stimulus repeats were available for analysis, yielding very reliable estimates. We defined the center frequency for a gamma rhythm as the frequency at which the change in power (in these averaged PSDs) was maximum within that gamma range. For SSVEP experiment, we analyzed only the counter-phasing condition. Hence, we took the power at 32 Hz (twice the counter-phasing frequency, i.e. 16 cps) for analysis.

We generated scalp maps using the topoplot.m function of EEGLAB toolbox (Delorme and Makeig, 2004, RRID:SCR_007292), modified to show each electrode as a colored disc, with color representing the change in power of slow gamma/fast gamma/SSVEP from baseline in decibels (dB).

We calculated slopes for rejecting noisy electrodes (as described in Artifact Rejection subsection) by fitting PSD across all analyzable repeats for each individual unipolar electrode with a power-law function as follows:

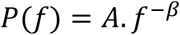

where *P* is the PSD across frequencies *f* ∈ [56 84] Hz. *A* (scaling factor) and *β* (slope) are free parameters obtained using least square minimization using the program *fminsearch* in MATLAB. We similarly estimated slopes for PSDs averaged across analyzable unipolar or bipolar electrodes during baseline period (−0.5 to 0 ms) for Supplementary Figure 1.

### Microsaccades and pupil data analysis

We detected microsaccades using a threshold-based method described earlier (Murty et al., 2018), initially proposed by (Engbert, 2006). In brief, we categorized eye movements with velocities that crossed a specified threshold for at least a specified duration of time as microsaccades. We set the velocity threshold between 3-6 times the standard deviation of eye-velocities and minimum microsaccade duration between 10-15 ms for every subject so as to maximize the correlation between peak velocity and amplitude of all microsaccades for that subject (also called a “main sequence”, see Engbert, 2006 for details), while maintaining the minimum microsaccade velocity at 10°/s and the microsaccade rate between 0.5/s and 3.0/s.

The above algorithm was applied for the analysis period of −0.5 s to 0.75 s of stimulus onset. After removing the microsaccade-containing repeats, there were 128.1±71.1 (mean±SD, minimum 5) repeats for elderly subjects (n=226, excluding 1 subject for whom eye-data could not be collected) for anterolateral electrodes reported in Figure 7c. Results did not change when we discarded 13 elderly subjects with less than 30 repeats without microsaccades from analysis (data not shown).

EyeLink 1000 system recorded pupil data in arbitrary units for every subject since pupil data cannot be calibrated for this tracker. Hence, instead of directly comparing time-series of pupil data, we used coefficient of variation (CV, ratio of standard deviation to mean) for every repeat as a measure of pupillary reactivity to stimulus of that repeat. This simple measure scales standard deviation of a distribution with respect to its mean. This allows comparison of variation in different distributions without getting affected by the mean of the distributions. We calculated CV for each analyzable trial separately and calculated mean CV across trials for every subject for comparison.

### Statistical analysis

Our findings were based mainly on PSD plots and we used appropriate statistical methods (Pearson correlation, linear regression and ANOVA) to confirm our interpretations. We used one-way (or two-way, as necessary) ANOVA to compare means of bar plots in Figures 4c, 4d, 6a and 8c, although non-parametric tests on medians instead of means using Kruskal-Wallis test (not reported) yielded qualitatively similar results. For two-way ANOVA, we considered age-group and gender as independent factors although including their interaction effect in the model yielded qualitatively similar results (not reported). We used Bonferroni correction for multiple tests/comparisons wherever necessary.

## Results

We recorded EEG from 236 elderly subjects aged 50-88 years and 47 subjects aged 20-48 years while presenting full-screen sinusoidal grating stimuli on a computer monitor. The stimuli differed in spatial frequency and orientation. Each stimulus was presented for 800 ms following a baseline period of 700 ms. The subjects were required to maintain fixation throughout the trial (containing 2-3 stimuli), while their eye-data was recorded for offline analysis. For analysis, we pooled the data across the four orientations (0°, 45°, 90°, 135°) for all subjects, as orientation selectivity was reported to be very low for humans for both slow and fast gamma bands (Murty et al., 2018). Similarly, we pooled across three spatial frequencies (1-4 cpd) which were reported to elicit salient gamma oscillations in humans (Murty et al., 2018).

Figure 1 shows the results of an example subject, a 53 years old female. Trial-averaged evoked potentials were plotted (Figure 1a) for electrodes P3, P1, P2, P4, PO3, POz, PO4, O1, Oz, O2 for unipolar reference (Figure 1a, left column) and PO3-P1, PO3-P3, POz-PO3, PO4-P2, PO4-P4, POz-PO4, Oz-POz, Oz-O1 and Oz-O2 (shown as dots in scalp maps in Figure 1c) for bipolar reference (Figure 1a, right column). These traces revealed a transient in the first 250 ms of stimulus onset and after the stimulus offset (i.e. after 800 ms). For the same set of electrodes, trial-wise power spectrograms were averaged to generate raw spectrogram and change in power spectrogram (w.r.t. a baseline period of −500 ms to 0 ms of stimulus onset), plotted in Figure 1a. Although not noticeable in the evoked potential traces and raw spectrograms, these stimuli elicited prominent gamma band responses as seen in the change in power spectrograms. These responses were in slow gamma (∼20-34 Hz) and fast gamma (∼36-66 Hz) range. Consistent with previous results (Murty et al., 2018), these responses were seen during the stimulus period (after the onset-transient) and were best noticed for bipolar reference (compared to unipolar reference). Also, slow gamma power showed a gradual build-up whereas fast gamma power showed a decreasing trend with stimulus duration (Figure 1a, bottom row). Alpha (8-12 Hz) power suppression was very weak in this subject. We also plotted power spectral densities (PSD) in the baseline period (dotted black trace in Figure 1b) and stimulus period (250 ms to 750 ms; solid black trace in Figure 1b) and change in power spectrum (blue trace in Figure 1b). Prominent ‘bumps’ in the slow and fast gamma range were noticeable in PSD in the stimulus period as well as change in spectrum. Also, no ‘bump’ was noticeable in the baseline PSD in the alpha range for this subject. These changes were most prominent in the parieto-occipital and occipital electrodes, as seen in the scalp maps for the bipolar reference case in Figure 1c.

**Figure 1.**
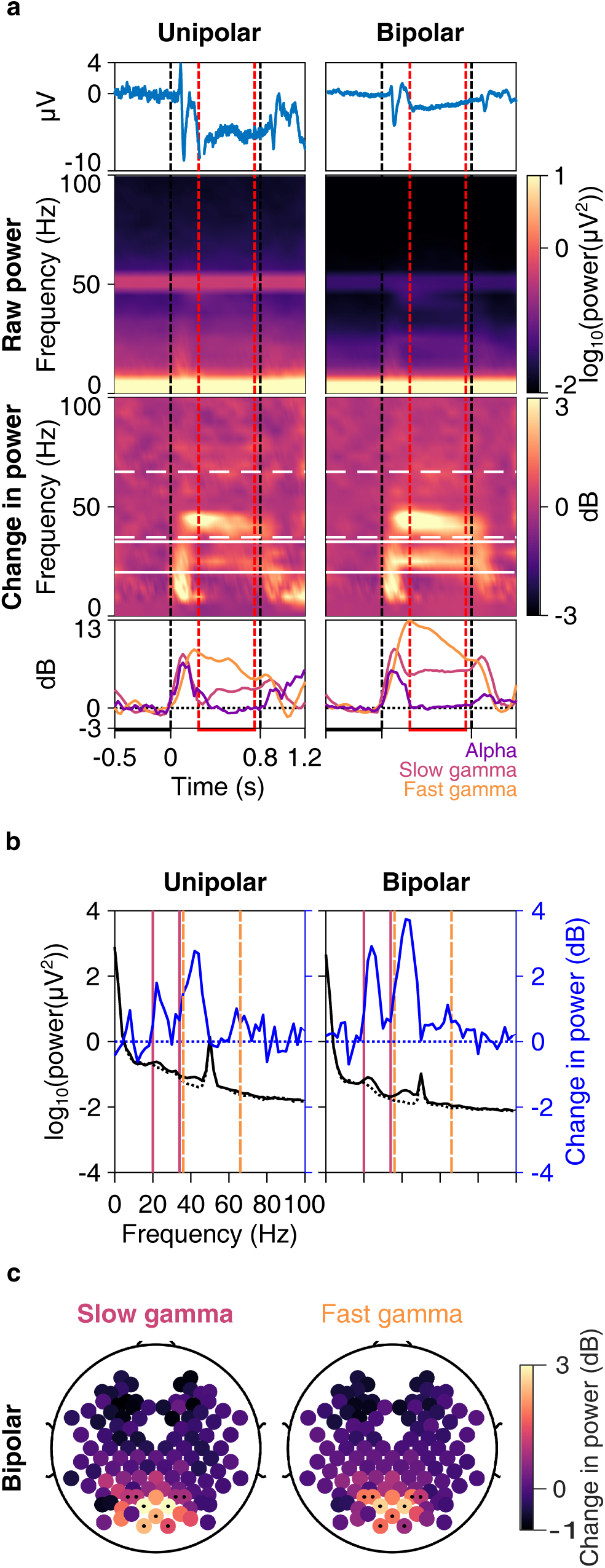
Slow and fast gamma in an example elderly subject. a) Trial-averaged EEG trace (1^st^ row, blue); time-frequency spectrograms of raw power (2^nd^ row) and change in power from baseline (3^rd^ row); and change in power with time (4^th^ row) in alpha (8-12 Hz, violet), slow (20-34 Hz, pink) and fast gamma (36-66 Hz, orange) bands averaged across 10 unipolar (left column) and 9 bipolar (right column) electrodes. Vertical dashed lines represent actual stimulus duration (0-0.8 s, black) and period used for analysis within stimulus duration (0.25-0.75 s, red). Horizontal lines represent baseline (−0.5-0 s, black) and stimulus (0.25-0.75 s, red) analysis periods. White lines in spectrograms represent slow (solid) and fast (dashed) gamma frequency ranges. b) Right ordinate shows raw power spectral densities (PSDs, black traces) vs frequency in baseline (dotted) and stimulus (solid) periods averaged across 10 unipolar electrodes (left column) and 9 bipolar (right column) electrodes; left ordinate shows the same for change in PSD (in dB, solid blue trace) in stimulus period from baseline. Solid pink lines and dashed orange lines represent slow and fast gamma bands respectively. c) Scalp maps showing 112 bipolar electrodes (represented as disks). Colour of each disk represents change in slow (left) and fast (right) gamma power. 9 electrodes used in 1a and 1b (right column) are marked with dots.

Our primary emphasis was to characterize gamma and other spectral signatures as a function of age within the elderly population (>49 years), for which we divided these subjects into two groups: 50-64 years (95 subjects; 51 female) and >64 years (141 subjects, 53 female). For completeness, we also show results from a cohort of younger subjects aged between 20-49 years (47 subjects; 16 female).

### Baseline power of slow and fast gamma did not vary with age

A recent study (Voytek et al., 2015) has suggested rotation of PSDs with age such that power at frequencies above ∼15 Hz is higher in elderly subjects compared to younger subjects, leading to flatter PSDs in elderly subjects. This could potentially bias the estimation of change in power in slow and fast gamma range in subjects of different age groups as higher baseline power in these rhythms in older subjects may lead to lower estimates of change in power. Hence, we first checked whether there was any difference in baseline PSDs across age. We calculated mean baseline PSDs of 10 unipolar electrodes and 9 bipolar electrodes separately, as mentioned above. We compared PSDs between 2-200 Hz in the two elderly groups as well as the younger group (Figures 2a and 2b for males and females), and males versus females (averaged across all ages; Figure 2c).

**Figure 2.**
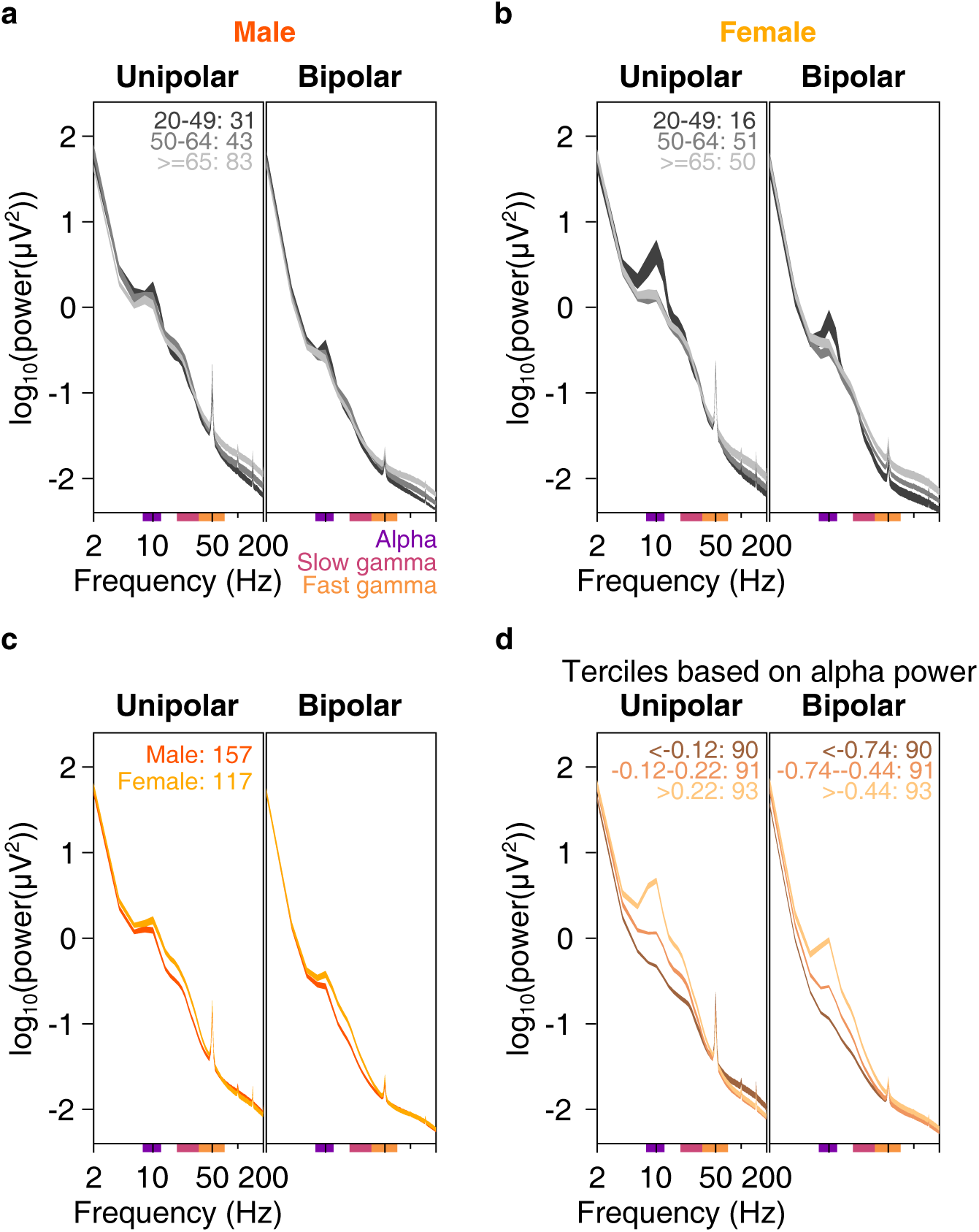
Baseline PSDs, slopes and alpha power. Baseline PSDs (averaged across 10 unipolar or 9 bipolar electrodes) for three age-groups on a log-log scale for unipolar (left) and bipolar (right) reference, plotted for males (2a) and females (2b). Thickness of traces indicate SEM across subjects. Age-group limits and the number of subjects in the respective age-groups are indicated on the left plot. c) Same as in 2a and 2b, but for males and females, pooled across all age-groups. d) Mean baseline PSDs for three ranges of baseline alpha power (8-12 Hz, power ranges for respective traces indicated on the plots) pooled across all age-groups. Thickness of traces and numbers indicate SEM across subjects and number of subjects in respective alpha power ranges. Coloured bars on the abscissa indicate alpha (8-12 Hz, violet), slow (20-34 Hz, pink) and fast gamma (36-66 Hz, orange) frequency bands.

Because our primary emphasis was comparison within the elderly group, we first compared the PSDs between the two elderly subgroups (dark and light gray traces in Figures 2a and 2b). The PSDs indeed appeared to become flatter with age (light gray trace was above the dark gray trace), but this effect was prominent only at frequencies above ∼50 Hz. In the slow and fast gamma ranges (indicated by coloured bars on the abscissa of plots in Figure 2), the two gray traces were largely overlapping. To quantify this, we performed a two-way ANOVA on baseline powers of alpha, slow gamma and fast gamma with age-group (50-64 or >64 years) and gender as factors and found that effect of age group was not significant for power in any band (p>0.05 in all cases except for fast gamma in bipolar case where p=0.03, which was not significant at Bonferroni corrected significance level of 0.05/3 = 0.016). Results were not qualitatively different when we performed one-way ANOVA for baseline power of alpha/slow/fast gamma across age-groups for males and females separately (p>0.05 for all cases except for fast gamma in females for bipolar case where p=0.03).

We obtained similar trends for comparisons (one-way ANOVA separately for males and females) between younger (<50 years) and elderly subjects (50 years and above). Baseline powers in alpha/slow/fast gamma ranges were not significantly different for younger and elderly male subjects in either reference schemes (Figure 2a, p>0.05 for all cases). However, elderly females had more baseline fast gamma power compared to younger females (*F*(1,115)=7.9, p=0.006) in bipolar case and lesser alpha power in both unipolar (*F*(1,115)=17.6, p=5.5*10^-5^) and bipolar (*F*(1,115)=6.5, p=0.012) cases (Figure 2b). These differences could be due to a small sample size of females in the younger age-group (n=16).

Across genders, females had significantly higher baseline slow gamma power than males (Figure 2c, data pooled across all 274 subjects; one-way ANOVA across gender: *F*(1,272)=24.5/27.9, p=1.3*10^-6^/2.6*10^-7^ for unipolar/bipolar reference schemes) and higher alpha power (*F*(1,272)=4.6/8.4, p=0.03/0.004 for unipolar/bipolar conditions). However, baseline fast gamma power was not significantly different (p>0.05 for both reference schemes). Amongst the elderly subjects (n=227, data not shown), females had only higher slow gamma compared to males (*F*(1,225)=16.4/21.3, p=7.1*10^-5^/6.4*10^-6^ for unipolar/bipolar conditions for slow gamma, *F*(1,225)=5.3, p=0.022 for alpha in bipolar case and p>0.05 for all other cases).

To test for the rotation of PSDs with age as suggested by Voytek *et al.* (2015), we computed the slopes between 16-44 Hz (Supplementary Figure 1; see Methods for details; this range was chosen to avoid the bump in the alpha band at the lower end and the 50 Hz noise at the higher end). Two-way ANOVA with age (young and elderly) and gender (male and female) as factors showed no significant difference in the slopes between young and elderly subjects for either unipolar or bipolar reference scheme case (p>0.05). However, females had steeper slopes compared to males (*F*(1,271)=7.9, p=0.005 and *F*(1,271)=31.4, p=5.1*10^-8^ for unipolar and bipolar cases respectively, Supplementary Figure 1a). Since females had higher baseline alpha power compared to males (Figures 2c), we tested whether any differences in baseline PSD slopes could be because of differences in baseline alpha power. We divided baseline PSDs of all subjects (young and elderly pooled together) into terciles based on alpha power (Figure 2d). Subjects who had higher baseline alpha power also had steeper PSD slopes. Regression of PSD slopes in 16-44 Hz frequency range with baseline alpha power was significant for both reference schemes (Supplementary Figure 1b). Further, when we performed partial correlation of slopes with age and baseline alpha power, slopes were significantly correlated with alpha power (rho=0.57, p=5.4*10^-25^ and rho=0.58, p=3.1*10^-26^ for unipolar and bipolar cases respectively) but not with age (rho=0.07 and −0.12 for unipolar and bipolar, p>0.05 for both). Thus, PSD slope was not influenced by age, but by baseline alpha power. We discuss these results in the context of the findings of Voytek and colleagues in the Discussion.

### Gamma was observed in more than 80% of subjects

As reported in our earlier study (Murty et al., 2018) and as in Figure 1, gamma was best observed, as a response to full-screen 100% contrast Cartesian visual gratings, in bipolar referencing scheme compared to unipolar. Hence, we limited further analysis to bipolar referencing. We divided the 9 bipolar electrodes mentioned above into 3 groups (Figure 3d): PO3-P1, PO3-P3, POz-PO3 (left anterolateral group); PO4-P2, PO4-P4, POz-PO4 (right anterolateral group) and Oz-POz, Oz-O1, Oz-O2 (posteromedial group). For each subject, we chose the electrode group that had maximum change in power in slow and fast gamma ranges added together. We labelled a subject as having either of the gamma rhythms if the change in power in these rhythms during stimulus period (calculated from data pooled across electrodes chosen for the subject) exceeded an arbitrarily chosen threshold of 0.5 dB from baseline. Figure 3a shows scatter plot of slow versus fast gamma change in power for all subjects. Based on our threshold, ∼84% of subjects had at least one gamma (slow: ∼77% and fast: ∼64%), while ∼57% of subjects had both the gammas, which could be observed as distinct “bumps” in the change in PSD from baseline (Figure 3b). Figure 3c shows the percentage of subjects in each age-group who had no/slow/fast/both gammas based on our threshold. The percentage of subjects who had only fast gamma or both gammas was highest in 20-49 years age-group and lowest in >64 years age-group.

**Figure 3.**
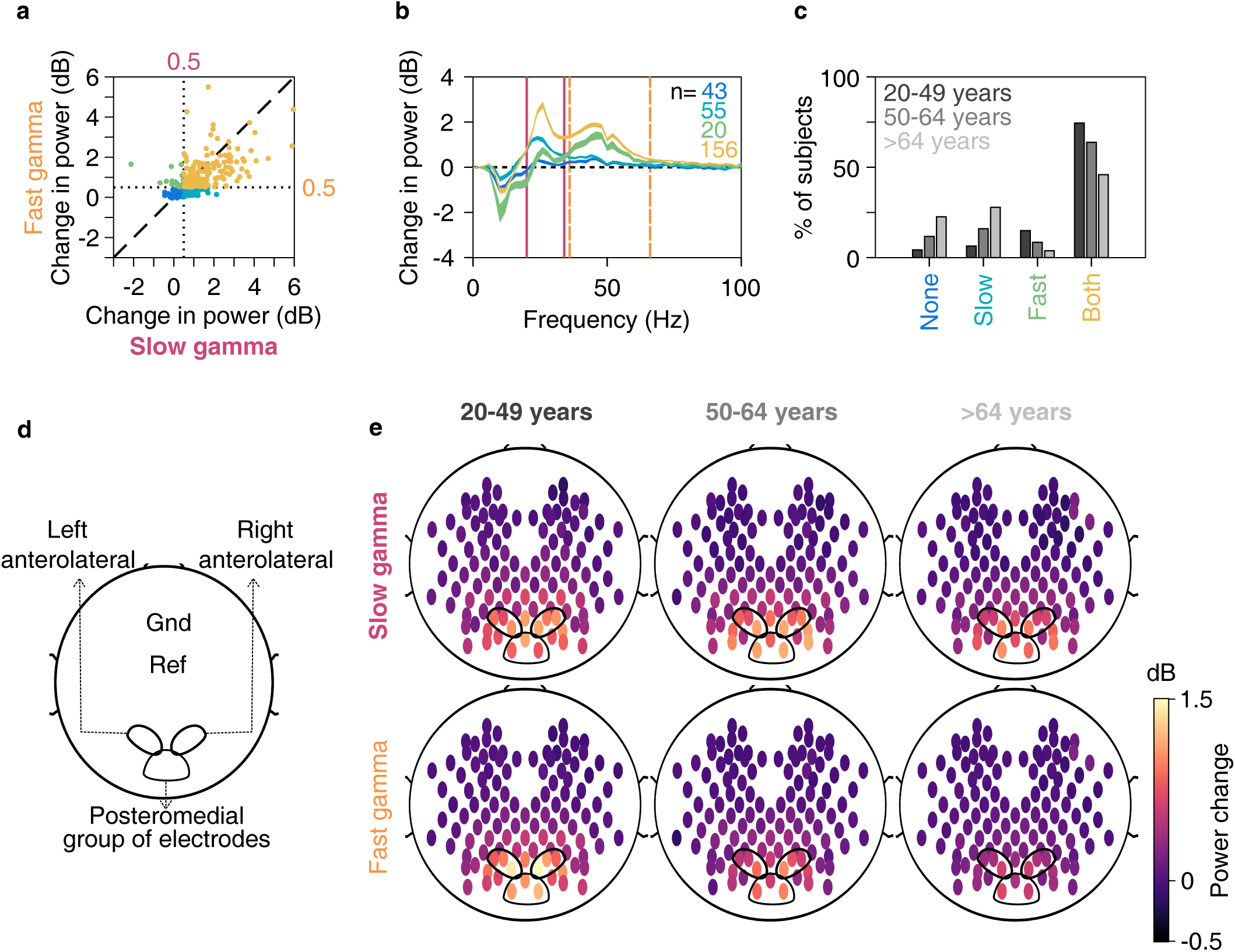
Slow and fast gamma in younger and elderly subjects. a) Scatter plot showing change in slow (abscissa) and fast (ordinate) gamma power. Dotted lines represent 0.5 dB threshold. Points represent subjects with no gamma (dark blue), only slow gamma (light blue), only fast gamma (green) and both gamma rhythms (yellow) with change in power above 0.5 dB threshold. b) Change in PSDs vs frequency averaged across subjects (numbers denoted by n) as categorized in 3a. Thickness of traces indicate SEM. Solid pink and dashed lines represent slow and fast gamma ranges respectively. c) Bar plot showing percentage of subjects in three age-groups (marked by respective colours) categorized as in 3a. d) Schematic showing placements of left and right anterolateral and posteromedial group of electrodes used for analysis on the scalp, as well as ground and reference electrodes. e) Average scalp maps of 112 bipolar electrodes (disks) for three age-groups for slow (top row) and fast (bottom row) gamma. Colour of disks represents change in respective gamma power. Electrode groups represented as in 3d.

Figure 3e shows change in power in slow (top row) and fast (bottom row) gamma rhythms across all electrodes (plotted as disks) for the young (left column) and the two elderly age-groups (middle and right columns). Both gamma rhythms were best observed in the same 9 bipolar electrodes mentioned above and depicted in Figures 3d and 3e. Further, power in both gamma bands appeared to decrease with age across the two elderly age-groups, although the results were more prominent for fast gamma.

### Change in gamma power was negatively correlated with age

To quantify this difference, we tested how gamma oscillations correlated with age in these electrode groups. We tested for anterolateral and posteromedial groups separately (Figures 4 and Supplementary Figure 2 respectively). For Figure 4, out of the left and right anterolateral groups, we chose that group which had maximum slow and fast gamma power change summed together. Figures 4a and 4b show mean change in spectrograms and PSDs respectively for the three age-groups separately for males and females. These plots highlight all the major results discussed later. First, both slow and fast gamma power reduced with age. This was observed between young and elderly groups (black versus the other two traces), and also within the two elderly sub-groups (dark and light gray traces). Second, peak frequencies of both slow and fast gamma reduced with age. Third, alpha suppression (change in 8-12 Hz power from baseline) in the stimulus period was more pronounced in young versus elderly, but there was no difference between the two elderly sub-groups.

**Figure 4.**
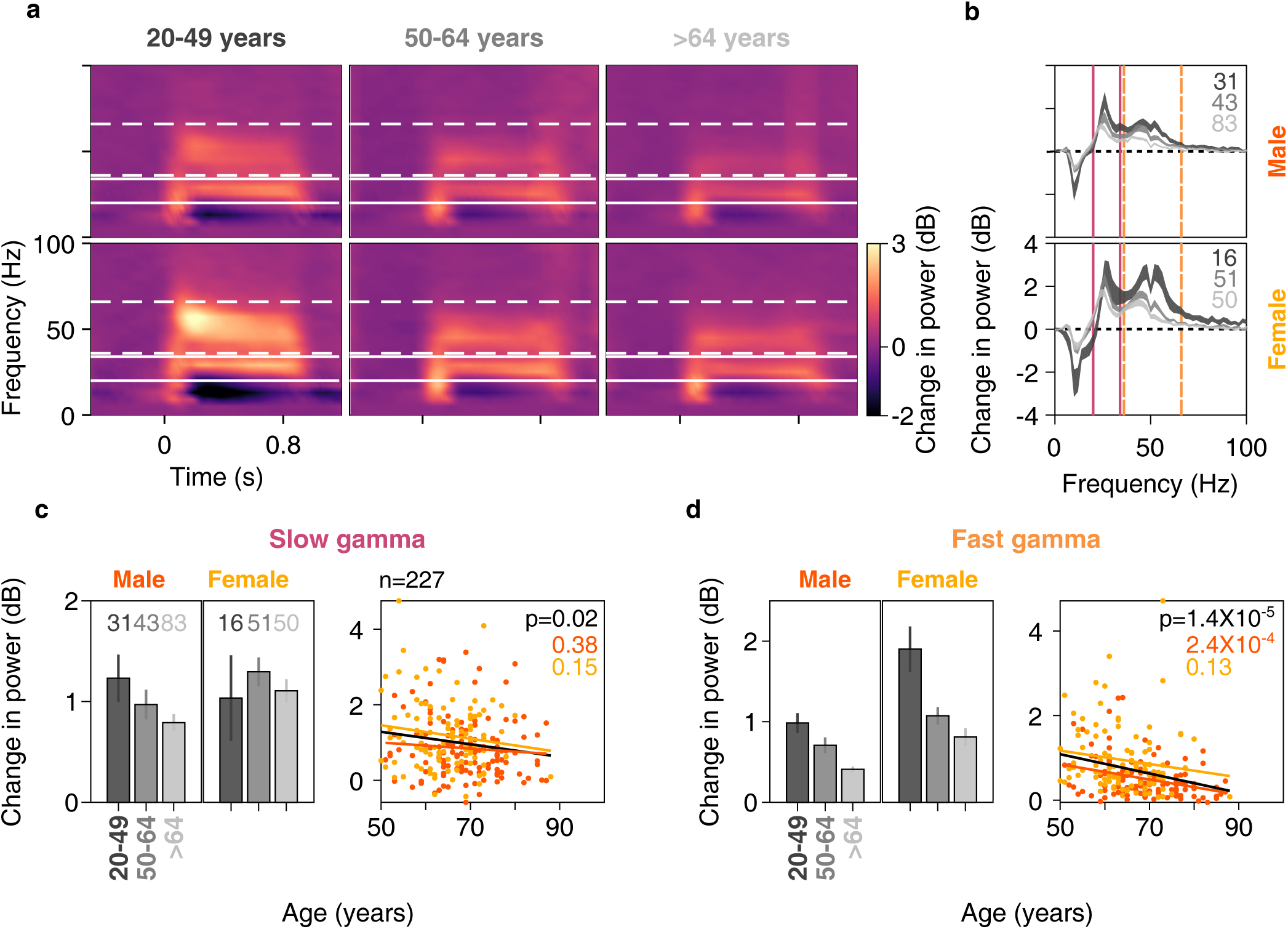
Change in gamma power vs age for anterolateral group of electrodes. Mean time-frequency change in power spectrograms (4a) and change in power spectra vs frequency (4b) for three age-groups separately for males (top row) and females (bottom row). Thickness of traces and numbers in 4b indicate SEM and number of subjects respectively. Solid and dashed lines indicate slow and fast gamma frequency ranges respectively. c) Left column: bar plots showing mean change in slow gamma power for three age-groups separately for males and females. Number of subjects for respective age-groups are indicated on top. Error bars indicate SEM. Right column: scatter plot for change in slow gamma power vs age for all elderly subjects (>49 years age-group, n=227), plotted separately for males (in orange) and females (in yellow). Orange, yellow and black solid lines indicate regression fits for males, females and data pooled across gender respectively. p-values of the regression fits are indicated in respective colours. d) Same as in 4c but for fast gamma.

The first observation was also reflected in the gamma power computed within the pre-specified ranges (as shown in the bar plots shown in Figure 4c and 4d), but there were some caveats. We computed the total power within a pre-specified band, which typically has larger contribution from lower frequencies because the absolute power is larger compared to higher frequencies within the band (which is not reflected in Figure 4b because it only shows the change in power). Consequently, if the traces are overlapping at lower frequencies within the band and diverge at higher frequencies, which was the case in the slow gamma range for both males and females (Figure 4b), the total power in the band may not be significantly different. In particular, for young females, the power at the start of the slow gamma band (20-26 Hz) was slightly lower than the elderly subgroups (Figure 4b, bottom plot, black versus gray traces), but became higher at higher frequencies within the slow gamma band (28-34 Hz). However, because the absolute power is higher between 20-26 Hz than 28-34 Hz, the total slow gamma power was actually lower for young females compared to elderly (Figure 4c, black versus gray bars). These issues can be partially addressed by changing the frequency range over gamma is computed (dependent on age and potentially even across subjects), but then the results are dependent on the level of customization of ranges, which we wanted to minimize. We observed that younger subjects had significantly more fast but not slow gamma than elderly subjects (two-way ANOVA with age-group (20-49 and >49 years) and gender as factors, *F*(1,271)=1.3/35.6, p=0.2/7.6*10^-9^ for slow/fast gamma across age-groups). Also, females had more slow and fast gamma than males (same two-way ANOVA, *F*(1,271)=4.7/37.9, p=0.03/2.5*10^-9^ for slow/fast gamma).

Among the elderly subjects, visual inspection of change in spectrograms and spectra revealed that both slow and fast gamma power was less in subjects of >64 years age-group compared to 50-64 years age-group. This trend was also noticeable in the bar plots in Figures 4c and 4d for both genders. As before, it was significant only for fast gamma (two-way ANOVA with age-group (50-64 and >64 years) and gender as factors; *F*(1,224)=2.4/11.4, p=0.12/8.4*10^-4^ for slow/fast gamma across age-group). Females had higher slow and fast gamma compared to males (same two-way ANOVA, *F*(1,224)=7.4/21.7, p=0.007/5.4*10^-6^ for slow/fast gamma across gender). We further quantified this observation by regressing change in slow and fast gamma power across age (scatter plots in Figures 4c and 4d). When the regression was done separately for males and females, the slopes were always negative (males: β=-0.008/-0.018 and females: β=-0.018/-0.016 for slow/fast gamma) but did not reach significance except for fast gamma in elderly males (p=2.4*10^-4^). When we pooled data across both genders, the results were significant (linear regression, β=-0.02, *R*^2^=0.02, p=0.022 and β=-0.02, *R*^2^=0.08, p=1.4*10^-5^ for slow and fast gamma respectively). Similar, albeit weaker results were observed in the posteromedial group of electrodes for slow gamma (Supplementary Figure 2c; linear regression, β=-0.016, *R*^2^=0.02, p=0.04) as well as fast gamma (Supplementary Figure 2d; β=-0.02, *R*^2^=0.04, p=0.001).

### Centre frequency of slow and fast gamma was negatively correlated with age

Gamma peak centre frequency was shown to decrease with age in an age group between 8-45 years (Muthukumaraswamy et al., 2010; Gaetz et al., 2012). This was observed in our data as well as noted above. To examine the change in centre frequency of slow and fast gamma rhythms in elderly in more detail, we plotted the change in power spectra (frequencies mentioned on abscissa) vs age (on ordinate, arranged in increasing order from top to bottom) of all 227 elderly subjects, separately for anterolateral (left column) and posteromedial (right column) group of electrodes. We defined centre frequency for each gamma as the frequency that had maximum change in power in the frequency range of that gamma, provided the total change in power in that gamma band was greater than our threshold of 0.5 dB (represented by circles and triangles for slow and fast gamma in Figure 5a; number of subjects having slow and fast gamma power change above this threshold is mentioned in Figure 5b). Figure 5b shows the same result as scatter plots of centre frequencies of slow (top row) and fast gamma (bottom row) plotted against the age of the subjects. Solid lines in Figures 5a and 5b indicate regression fits of centre frequencies against age (showing decreasing trend). This trend was significant for both slow and fast gamma in the anterolateral group (linear regression for centre frequency vs age: β=-0.08, *R*^2^=0.04, p=0.008 for slow gamma and β=-0.16, *R*^2^=0.06, p=0.008 for fast gamma). Similar, albeit weaker results were observed for the posteromedial group (fast gamma: β=-0.17, *R*^2^=0.06, p=0.008; slow gamma: β=-0.06, *R*^2^=0.02, p=0.052). Note that because our analysis was done over 500 ms of data, the frequency resolution was 2 Hz, which limited our ability to observe small shifts in the peak frequency.

**Figure 5.**
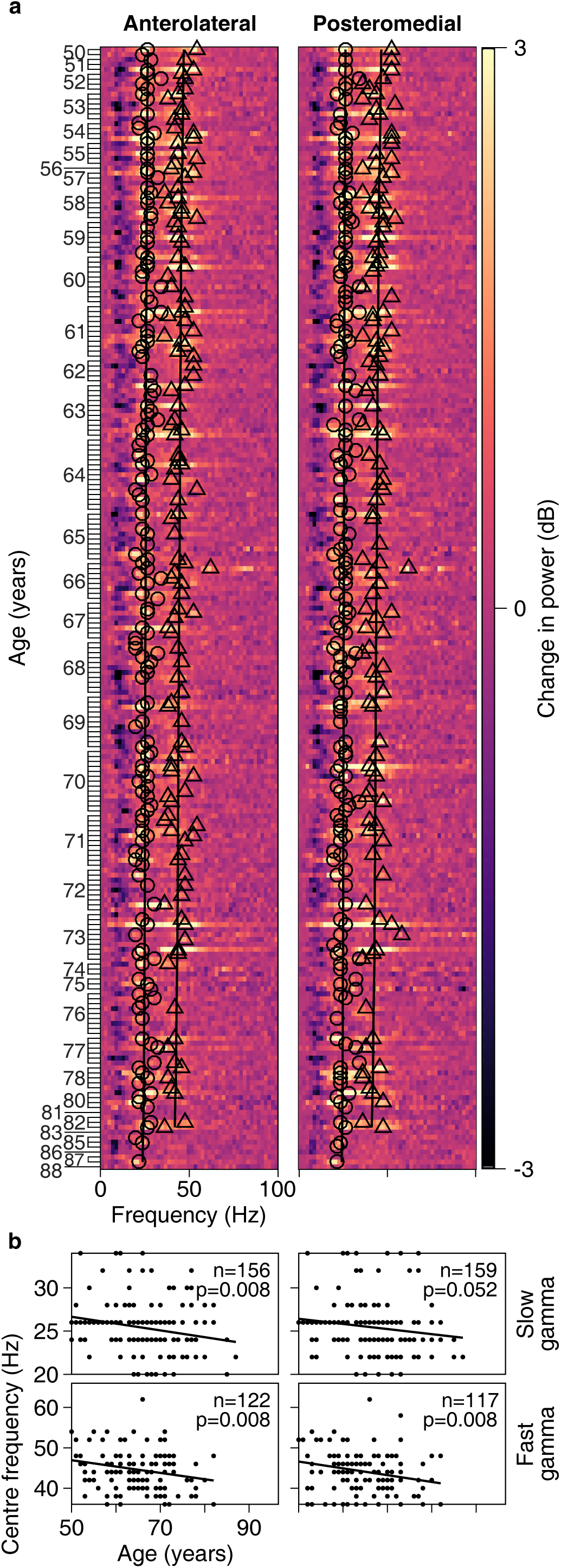
Centre frequency of slow and fast gamma vs age for elderly subjects. a) Change in power vs frequency for 227 elderly subjects arranged in ascending order of age. Age for each subject is indicated on ordinate. Colour represents change in power in dB for each frequency. Circles and triangles represent centre frequency of slow and fast gamma rhythms respectively, indicated only for those subjects who have change in power in respective gamma range above 0.5 dB. b) Scatter plots showing centre frequency vs age for slow (top row) and fast (bottom row) gamma for those subjects who have change in power in respective gamma range above 0.5 dB (numbers indicated on the plots). Solid lines in both 5a and 5b are regression fits for centre frequency vs age. p-values for these fits are indicated on plots in 5b. Left and right columns show analysis for anterolateral and posteromedial group of electrodes respectively.

### Alpha suppression did not reduce with age in elderly subjects

We noticed prominent alpha suppression for younger as well as elderly subjects, as noted above (and in Supplementary Figures 2a and 2b). Alpha suppression was stronger in younger subjects compared to elderly subjects (data for anterolateral group is shown in Figure 6a; two-way ANOVA with age-groups (20-49 and >49 years) and gender as factors; *F*(1,271)=33.2, p=2.2*10^-8^ across age-groups), but did not differ significantly between genders (*F*(1,271)=0.5, p=0.49 across gender). Amongst elderly subjects however, alpha suppression did not differ across age-groups (50-64 and >64 years) and gender (two-way ANOVA, p>0.05 for both age-group and gender). We further confirmed this observation by regressing alpha suppression across age for all the elderly subjects (scatter plot in Figure 6a). Alpha suppression was not significantly correlated with age for either gender or for data pooled across genders (p>0.05 for all cases). The trends were not qualitatively different when we repeated the analysis for the posteromedial group of electrodes. This is also observed in the scalp maps shown in Figure 6b. We did not test for alpha frequency as 2 Hz frequency resolution of our PSDs left us with only 3 frequency points in the alpha range (8, 10 and 12 Hz).

**Figure 6.**
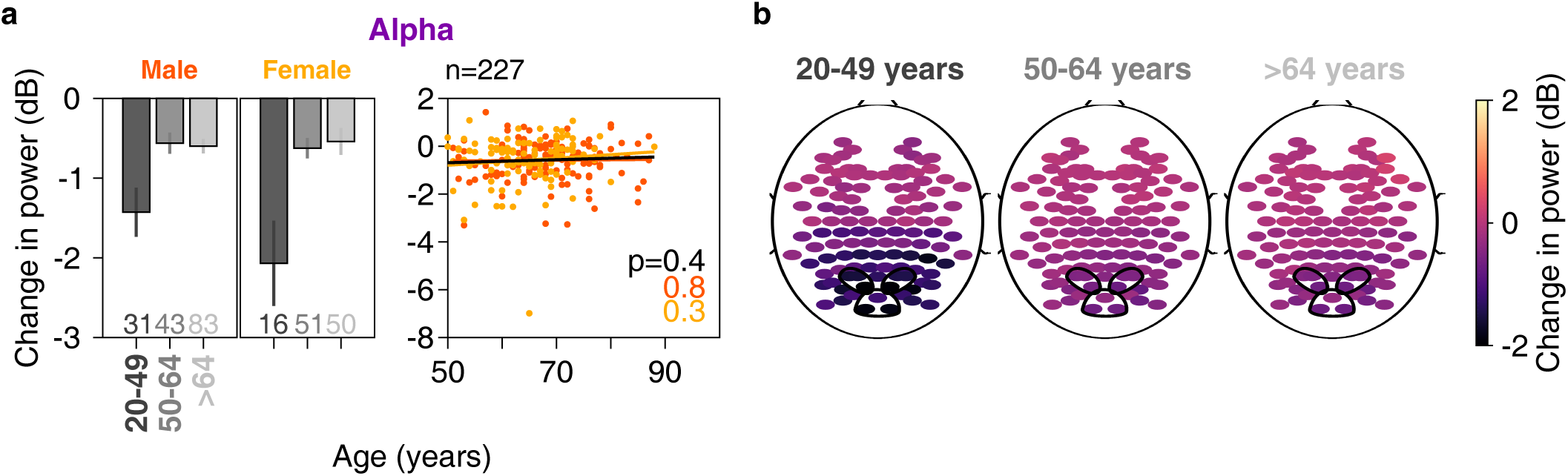
Change in alpha power vs age. a) Left column: bar plots showing mean change in alpha power across anterolateral group of electrodes for three age-groups separately for males and females. Number of subjects for respective age-groups are indicated at bottom. Right column: scatter plot for change in alpha power vs age for all elderly subjects (>49 years age-group, n=227), plotted separately for males (in orange) and females (in yellow). Same format as in Figure 4c. b) Scalp maps for 112 electrodes (disks) averaged across all subjects separately for three age-groups. Colour indicates change in alpha power for each electrode, same format as in Figure 3e.

### Microsaccades and pupillary reactivity did not contribute to negative correlation between change in gamma power and age

Next, we studied the potential contribution of eye-movement (including microsaccades) and pupillary diameter on our results. Figure 7a shows mean eye-position for each of the elderly age-groups in horizontal (top row) and vertical (middle row) directions (n=226, eye data was unavailable in one subject; thickness represent SEM). Eye-position did not vary in the two age-groups in either direction. Further, we extracted microsaccades for every analysed trial for every subject in the two age-groups (see Materials and Methods), yielding a total of 27350 and 38373 microsaccades for the two age-groups. Figure 7b shows a scatter plot of peak velocity versus maximum displacement for each microsaccade (a plot called “main sequence”, see Engbert, 2006). These microsaccade clouds were highly overlapping for these two groups and had similar microsaccade rates (0.80±0.05/s and 0.88±0.05/s). Histograms of microsaccade rate during −0.5 – 0.75 s of stimulus onset for both the elderly age-groups were also highly overlapping (Figure 7a, bottom row), although we see a trend of slightly higher microsaccade rate for subjects aged >64 years compared to 50-64 years age-group. We then computed power after removing trials containing microsaccades (see Materials and Methods for details), and could replicate the results in Figure 4: change in both slow and fast gamma power decreased with age significantly (β=-0.02, *R*^2^=0.03, p=0.015 for slow gamma and β=-0.02, *R*^2^=0.08, p=2.7*10^-5^ for fast gamma, Figure 7c).

**Figure 7.**
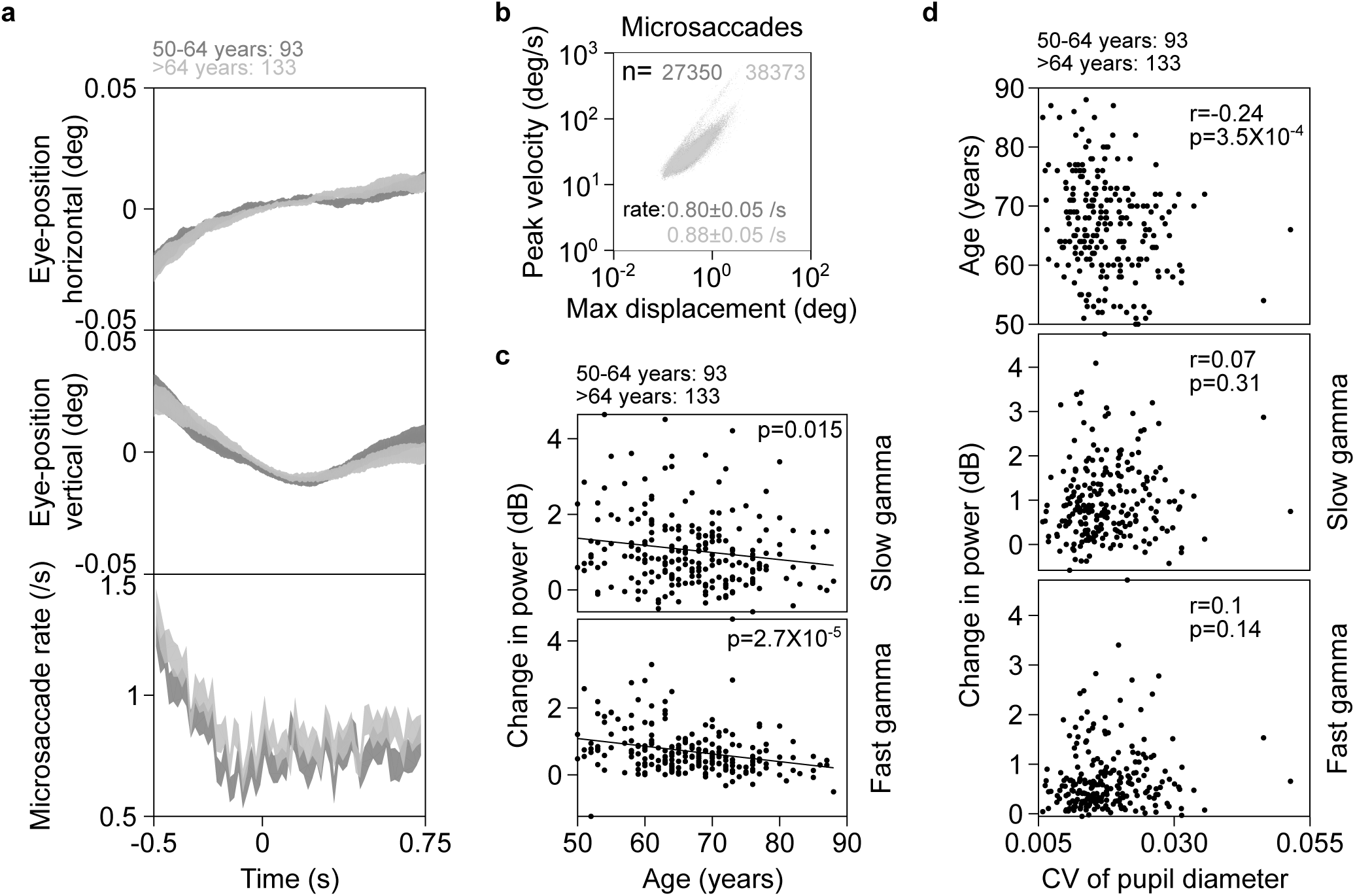
Eye position, microsaccades and pupillary reactivity across age for elderly subjects. a) Eye-position in horizontal (top row) and vertical (middle row) directions; and histogram showing microsaccade rate (bottom row) vs time (−0.5-0.75 s of stimulus onset) for elderly subjects (n=226). Number of subjects in each age-group is indicated on top. Thickness indicates SEM. b) Main sequence showing peak velocity and maximum displacement of all microsaccades (number indicated by n) extracted for both elderly age-groups. Average microsaccade rate (mean±SEM) across all subjects for each elderly age-group is also indicated. c) Scatter plot showing change in power vs age for slow (top row) and fast (bottom row) gamma for all elderly subjects with analysable data after removal of trials containing microsaccades. Solid lines indicate regression fits. Numbers of subjects with analysable data in each age-group is indicated on top. d) Scatter plots for coefficient of variation (CV) of pupil diameter vs age (top row), change in slow (middle row) and fast (bottom row) gamma power. Pearson correlation coefficients (r) and p-values are also indicated.

We next tested if pupillary reactivity to stimulus presentation affected change in gamma power with age. We calculated mean coefficient of variation (CV) of pupil diameter across every analysable trial for all 226 subjects (eye data was unavailable in one subject). We observed that mean CV decreased significantly with age in the elderly subjects, possibly because of senile miosis (Pearson correlation, r=-0.24, p=3.5*10^-4^, Figure 7d top row). However, neither slow nor fast gamma power varied with mean CV of pupil diameter (slow/fast: r=0.07/0.1, p=0.31/0.14 and r=0.09/0.12, p=0.19/0.06 for anterolateral (Figure 7d middle and bottom rows) and posteromedial electrodes respectively (data not shown)).

### 32 Hz SSVEP power was negatively correlated with age

Finally, we checked whether SSVEPs in the gamma range were affected by healthy aging. Specifically, we tested 32-Hz SSVEPs elicited by gratings counter-phasing at 16 cps. Figure 8a and 8b show change in power spectrograms and spectra respectively for males and females separately for the two elderly and the younger age-groups for the anterolateral group of electrodes, with same conventions as in Figures 4. We saw clear peaks at 32 Hz in both change in power spectrograms and PSDs. Insets in Figure 8b show a zoomed-in image of the respective change in PSDs to show the difference in these peaks for the three age-groups. Amongst the elderly age-groups, the mean SSVEP change in power was less in the >64 years age-group compared to 50-64 years age-group in both males and females. We regressed the SSVEP power change with age (scatter plot in Figure 8c bottom row, shown separately for males and females). Change in SSVEP power at 32 Hz decreased significantly with age for both males and females separately (males: β=-0.17, *R*^2^=0.09, p=0.002 and females: β=-0.18, *R*^2^=0.08, p=0.007) as well as when the data were pooled across genders (β=-0.19, *R*^2^=0.11, p=1.4 × 10^-6^, regression fit indicated by black line in bottom row of Figure 8c).

**Figure 8.**
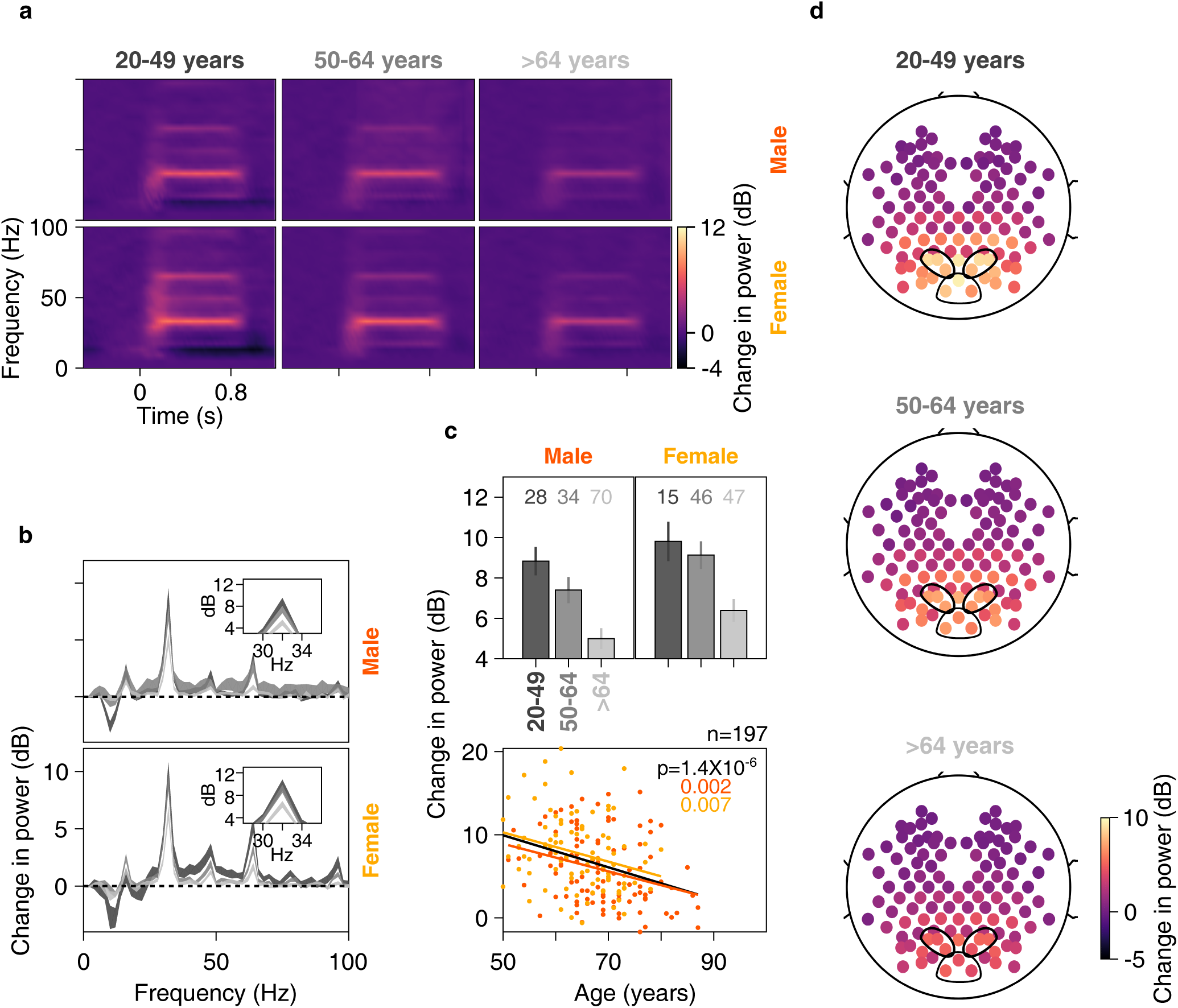
Change in SSVEP power vs age for posteromedial group of electrodes. Time-frequency change in power spectrograms (8a) and change in power spectra vs frequency (8b) for three age-groups separately for males (top row) and females (bottom row). Thickness of traces in 8b indicates SEM. Insets in 8b display zoomed-in images of respective main plots, showing clear SSVEP peaks at 32 Hz. c) Top row: bar plots showing mean change in SSVEP power for three age-groups separately for males and females; Numbers of subjects in each age-group is indicated on top. Error bars indicate SEM. Bottom row: scatter plot for change in SSVEP power vs age for all elderly subjects (>49 years age-group, n=197), plotted separately for males (in orange) and females (in yellow). Orange, yellow and black solid lines indicate regression fits for males, females and data pooled across gender respectively. p-values of the regression fits are indicated in respective colours. d) Scalp maps for 112 electrodes (disks) averaged across all subjects separately for three age-groups. Colour indicates change in SSVEP power at 32 Hz for each electrode.

We repeated this analysis for posteromedial group of electrodes and noticed similar results (regression of change in 32 Hz SSVEP power versus age for males: β=-0.16, *R*^2^=0.11, p=0.0008, females: β=-0.14, *R*^2^=0.06, p=0.02 and for data pooled across gender: β=-0.17, *R*^2^=0.11, p=1.5 × 10^-6^). We noticed this decrease of 32 Hz SSVEP power with age also in the mean scalp maps for all analysable electrodes across all subjects in the three age-groups, as depicted in Figure 8d.

## Discussion

We tested for age-dependent variation of stimulus-induced change in power and centre frequency of narrow-band gamma oscillations in both slow and fast gamma frequency ranges in healthy elderly subjects aged 50-88 years. We observed a decrease in power of both slow and fast gamma oscillations with age, although the decrease in fast gamma was more salient than slow gamma. On the other hand, level of alpha suppression did not change with age in elderly subjects. Finally, centre frequency of both gamma rhythms decreased with age in these subjects. As there was no significant change in baseline slow/fast gamma power, eye-position and microsaccade rate across age, we ruled out the possibility that the age-related variations in gamma could be because of such factors. Further, we also studied variation of 32 Hz SSVEP power with age and observed a negative correlation. All these results were also analysed for a cohort of younger subjects (aged 20-48), and the results were consistent with previous reports on gamma oscillations and some of the trends observed in the elderly population.

### Baseline alpha power and alpha suppression

Previous studies have suggested reduction in baseline alpha power in elderly subjects compared to younger subjects (Babiloni et al., 2006). Also, task-related modulation of alpha power was seen to be reduced in older adults compared to younger subjects (Vaden et al., 2012). Our results were similar to these previous reports: baseline alpha power was significantly higher in younger females versus elderly (Figure 2b), and showed a decreasing trend with age in males (although not significant, Figure 2a). Similarly, stimulus-induced alpha suppression was stronger for younger subjects compared to elderly subjects (Figure 6a). This is notwithstanding the different recording paradigms from previous studies: in our study, baseline alpha was recorded during eyes-open state (as opposed to resting, eyes-closed state in Babiloni and colleagues, (2006)) and alpha suppression was measured during passive fixation (as opposed to an active memory task in Vaden and colleagues (2012)). Among the elderly subjects, however, neither baseline alpha power (Figures 2a and 2b) nor alpha suppression (Figure 6a) varied with age. Different results for alpha versus gamma power (which decreased with age) in elderly subjects suggest different biophysical mechanisms of these oscillations.

### Baseline PSD slopes

Some authors have suggested that power-law distribution (1/*f*^β^, where β is the PSD slope) of brain electrical activity represents broadband scale-free activity of brain that is dependent on behavioral states (He et al., 2010; He, 2014; Podvalny et al., 2015) and cognitive abilities (Voytek et al., 2015; Sheehan et al., 2018). Specifically, Voytek and colleagues had suggested that flattening of PSD slopes might be a hallmark of senile physiological cognitive decline. In our study however, we did not notice any significant correlation between baseline PSD slopes and age in the unipolar reference scheme (as used by Voytek and colleagues), especially for elderly subjects. There are several reasons that could have led to this discrepancy. First, we estimated broadband slopes in the range of 16-44 Hz as opposed to 2-24 Hz (as in Voytek et al.). This is to avoid the contribution of baseline alpha power (8-12 Hz), against which we were testing for slopes (Figure 2d and Supplementary Figure 1b). Second, the sample size of Voytek and colleagues was small (11 young and 13 elderly) with a larger proportion of females in the younger group (male:female = 4:7 and 8:5 in young and elderly groups). Because females had steeper slopes than males (Figure 2c), underrepresentation of females in the elderly group could have led to flatter PSDs in their data. Finally, we found that PSD slopes were correlated with baseline alpha power (which was higher in younger versus elderly), but there was no dependence of slope on age when controlled for baseline alpha power (using partial correlation). Note that a similar correlation of slopes with alpha power in human MEG and EEG as well as monkey ECoG has also been reported by Muthukumaraswamy and Liley (2018).

We note, however, that the PSDs did tend to become flatter with age, albeit at a higher frequency range (>50 Hz; Figure 2a and 2b), consistent with the ECoG results of Voytek and colleagues and consistent with the neural noise hypothesis proposed by them. Further, our “spontaneous activity” used for PSD computation was during the fixation task itself, and PSDs were computed using segments of 500 ms, much less than the 2 second segments used by Voytek and colleagues. Consequently, the frequency resolution was 4 times higher in the study of Voytek and colleagues, which could have led to the identification of small changes in slopes better than ours. Longer stimulus-free epochs (at least 2 seconds or more), preferentially in both eyes closed and eyes open conditions are required to test whether the flattening of PSD slope occurs at lower frequencies as well.

### Possible confounds from ocular factors

Broadband induced gamma responses have been proposed to be correlated with occurrence of microsaccades (Yuval-Greenberg et al., 2008). However, in our previous study, we did not note any effect of microsaccades on orientation tuning of narrow-band slow and fast gamma oscillations in macaques (Murty et al., 2018). Consistently, we did not find any effect of microsaccades on age-dependent decrease of slow and fast gamma power in this study.

It is possible that retinal illuminance is reduced due to senile pupillary miosis, which is indirectly reflected in the reduced pupillary reactivity to stimulus presentation across age (Figure 7c). Other abnormalities of peripheral visual system like age-related increase in density of crystalline lens, age-related macular degeneration, etc. could have had affected our results (Owsley, 2011). The subjects did not undergo a thorough ophthalmic examination due to time limitations. However, we argue that the results presented here are likely due to neurophysiological effects of aging on two grounds. First, in addition to a reduction in gamma power, there is a reduction in gamma centre frequency with age, which is harder to explain based on the abnormalities listed above. Second, slow/fast gamma power was not dependent on pupillary reactivity to stimulus (Figure 7d). Nonetheless, we observed that the percentage of variance in the gamma power/frequency or SSVEP power explained by age is very less (maximum *R*^2^ among all cases was 0.11 for decrease in SSVEP power across age in posteromedial electrodes). Hence, we recognise that age is one of the many possible factors that influence gamma power/frequency and do not completely rule out the possibility that any hidden physiological variables could have had contributed to this variance.

### Possible mechanisms of age-related reductions in gamma frequency and change in power

Previous studies in MEG had reported significant positive correlations between (fast) gamma frequency and cortical thickness as well as volume of cuneus (Gaetz et al., 2012) and thickness of pericalcarine area (Muthukumaraswamy et al., 2010), measured through structural MRI. Further, age-related decreases in cortical volume, thickness and/or surface area were observed in various regions of the brain like precuneus, cuneus, lingual, pericalcarine and lateral occipital areas of the occipital cortex (Salat et al., 2004; Lemaitre et al., 2012). In line with these observations, these MEG studies have also noted decrease of centre frequency of gamma with age, esp. in young and middle-aged adults. Similarly, (fast) gamma peak frequency has been positively correlated with brain GABA levels (Edden et al., 2009; Muthukumaraswamy et al., 2009). Although we have not tested directly for these variables in our study, we speculate that these morphological and neurochemical factors are the reasons for our observations regarding fast gamma frequency. However, the factors behind reduction of slow/fast gamma power and slow gamma frequency with age remain open questions.

To conclude, our study throws light on various features of baseline spectra (like baseline alpha power and its relation to PSD slopes) and spectral responses to Cartesian gratings (alpha suppression, slow and fast gamma) in a large cohort of healthy elderly. Our study could thus act as normative for future gamma and SSVEP studies in the elderly age-group. Further, based on observations in previous rodent studies (for example, Verret et al., 2012; Iaccarino et al., 2016) as described before, some authors have suggested a causative role of (fast) gamma disruption in neurodegenerative disorders of aging such as AD (Palop and Mucke, 2016). Alternatively, our results suggest that gamma and SSVEPs suffer reduction in power with age even in the absence of cognitive decline. These studies taken together, decrease in gamma/SSVEP power may represent a continuum of healthy aging – preclinical cognitive decline – dementia spectrum and may act as a harbinger to senile or pathological cognitive decline, a hypothesis that needs to be tested in future studies.

---

**Supplementary Figure 1.**
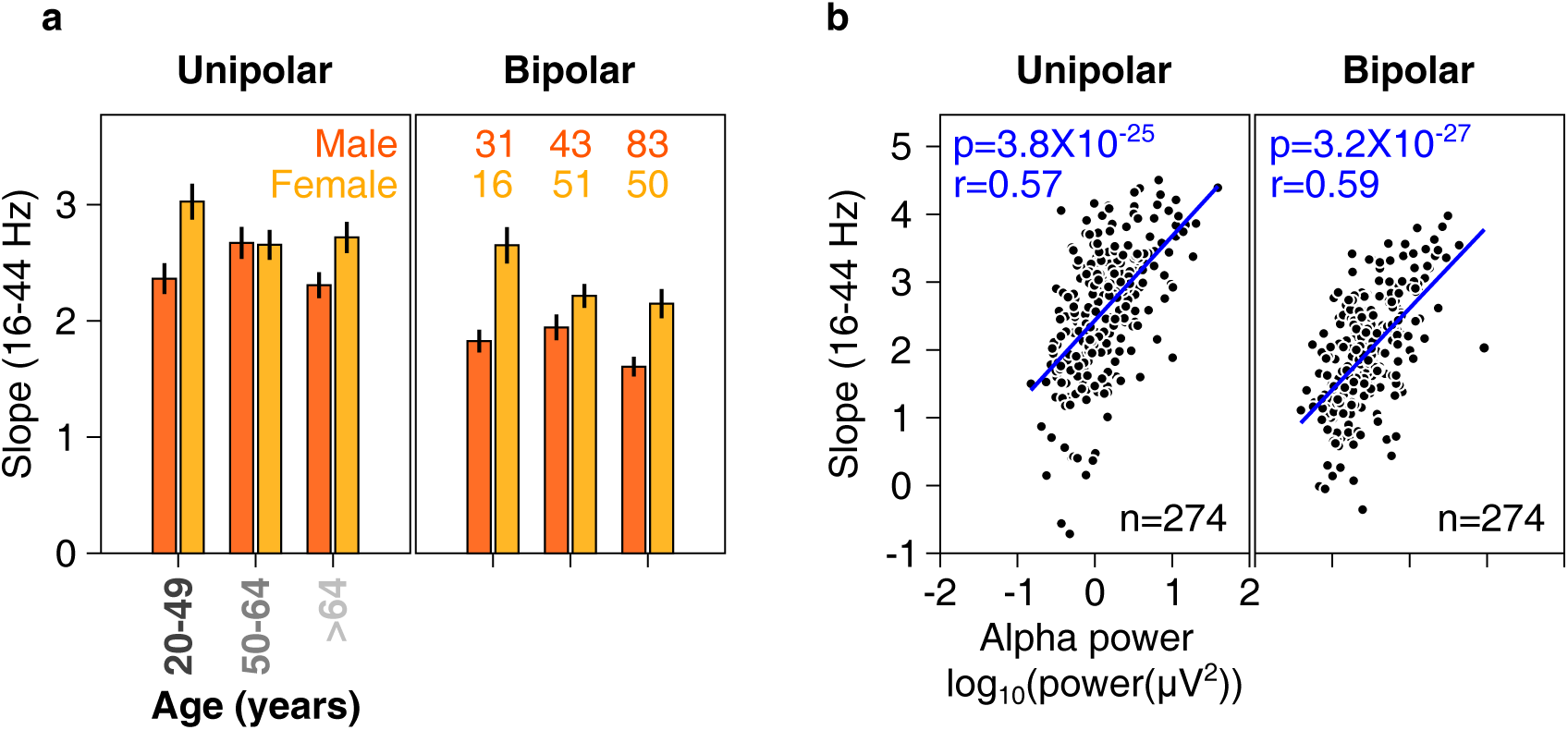
PSD slopes (16-44 Hz) across age-groups, gender and baseline alpha power. a) Bar plots showing mean slopes of baseline PSDs for three age-groups separately for males and females in 16-44 Hz range for unipolar (left) and bipolar (right) reference. Error bars indicate SEM. Numbers on top indicate number of subjects. b) Scatter plot showing baseline slopes in 16-44 Hz range vs alpha power, for unipolar (left) and bipolar (right) reference for all 274 subjects used for analysis (227 elderly and 47 younger). Lines indicate regression fits. Pearson correlation coefficients (r) and p-values are also indicated.

**Supplementary Figure 2.**
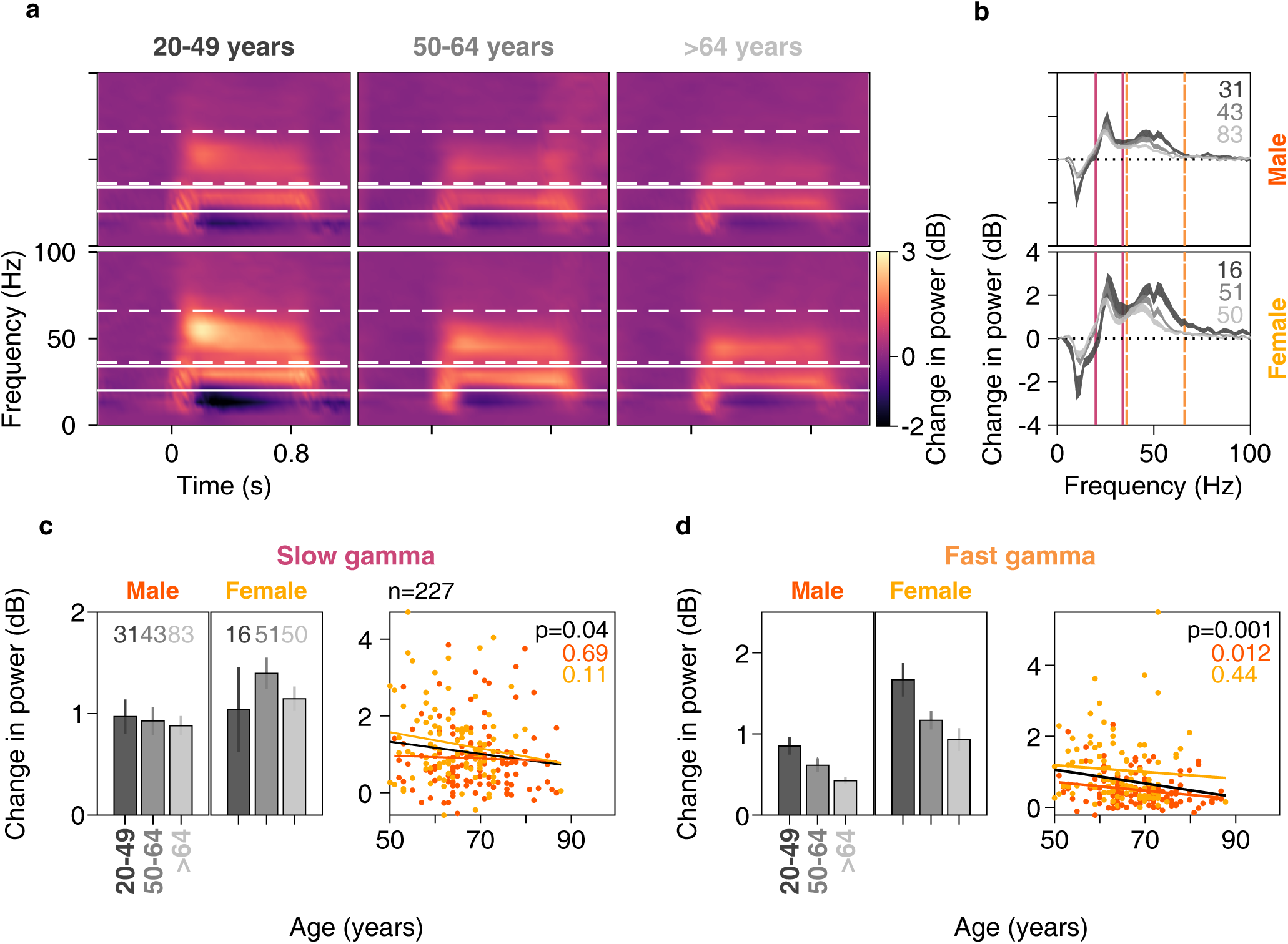
Change in gamma power vs age for posteromedial group of electrodes. Same format as in Figure 4.

